# The Hyaluronan Synthase isoforms direct distinct patterns of fibroblast activation linked to fibrosis progression versus resolution following renal ischaemia

**DOI:** 10.1101/2024.03.27.586996

**Authors:** Irina Grigorieva, Adam Midgley, Charlotte Brown, Gilda Chevez, Aled Williams, Dominic Manetta-Jones, Timothy Bowen, Deva Chan, Anna Plaas, Rafael Chevez, Robert Steadman, Usman Khalid, Soma Meran

**Author notes:** **Corresponding author:** Dr Soma Meran, Clinical Reader, Wales Kidney Research, Unit University Hospital of Wales, Heath Park, Cardiff. CF14 4XN. These authors contributed equally to this work and are joint first authors (Irina Grigorieva, Adam Midgley).

## Abstract

**Background:** The stroma plays a key role during renal development and in regeneration after injury. However, following injury, the stroma expands driving progressive fibrosis. Hyaluronan (HA) is a glycosaminoglycan that is absent in healthy kidney stroma but highly expressed in disease. To understand strategies to modulate stromal HA towards therapeutic advantage, this study compares HA and HA Synthase (HAS) enzyme expression in kidney development, health, disease, and recovery.

**Methods:** Rats underwent ischaemia reperfusion injury (IRI) with/without ischaemic preconditioning (IPC) and kidneys histologically analysed. Kidneys from C57BL/6 embryos and HAS1/3^-/-^ mice were also analysed and parallel mechanistic cell studies performed using primary human fibroblasts.

**Results:** In health, stromal HA was absent from the renal cortex. HAS1 was expressed in some epithelial cells, whilst HAS2 was not expressed. Following IRI there was increased stromal HA in areas of chronic fibrosis, alongside increased HAS2 (but not HAS1) expression. In contrast, during development prominent stromal HA matrices were evident in areas of tubular generation, with strong HAS1 (not HAS2) expression. Following IPC+IRI, stromal HA and HAS2 were attenuated; whilst HAS1^+^ cells expanded but were distinct from α-SMA^+^ myofibroblasts. Cell studies demonstrated that HAS1^+^ fibroblasts had a functionally distinct phenotype, with enhanced migration and FAP expression but attenuated α-SMA, EDA-FN and COL1A1 expression, whereas HAS2^+^ fibroblasts demonstrated a classic α-SMA^+^ contractile myofibroblast phenotype with high EDA-FN and COL1A1.

**Conclusions:** HA is a key regulator of stromal fibroblast heterogeneity, with HAS1 and HAS2 defining phenotypically distinct populations that may influence divergent renal outcomes following injury.

**SIGNIFICANCE STATEMENT:** Hyaluronan (HA) is a matrix glycosaminoglycans that is absent in healthy kidney cortex but demonstrates increased expression in the renal stroma during progressive fibrosis. This study makes comparisons of HA accumulation, localisation, and HA Synthase (HAS) protein expression during kidney development, in health, following ischaemic kidney injury and during renal recovery. The study identifies that different HAS isoenzymes (HAS1 and HAS2) mediate distinct functional fibroblast phenotypes *in vitro* and are localised in distinct stromal localisations and cell sub-populations *in vivo*. The data provides interesting insights into HA dependent regulation of fibroblast stromal heterogeneity and identifies the *novel* finding that HAS1 defines cell populations that are associated with kidney recovery following ischaemic injury and are protective against progressive renal fibrosis.

## INTRODUCTION

Over the last decade there has been increased appreciation that acute kidney injury (AKI) and chronic kidney disease (CKD) are not two completely distinct entities and that even minor episodes of AKI can lead to an increased burden of CKD^1–4^. The term ‘AKI-to-CKD continuum’ is therefore used to describe the pathophysiological association between AKI and CKD and these insights have led to studies that investigate the relationship between experimental AKI and progressive fibrosis. Ischaemia-reperfusion injury (IRI) is the leading cause of AKI in both native and transplanted kidneys^5, 6^. IRI describes the tissue damage that occurs when the blood supply to an organ is interrupted, followed by re-instatement of blood flow and subsequent re-oxygenation. Ischaemia results in tissue hypoxia, whilst the reperfusion paradoxically augments tissue injury through activation of immune, inflammatory and apoptotic pathways^7, 8^. In native kidneys IRI develops in the context of disease processes linked with poor renal perfusion such as sepsis, heart disease and shock. It affects between 8-15% of hospitalized patients and is associated with poor patient outcomes^3, 8, 9^. In transplanted kidneys, IRI is an inevitable consequence of the surgical process and results in delayed graft function, the severity of which influences eventual transplant outcomes. Identifying mechanisms that can alleviate kidney damage associated with IRI is therefore key to reducing the burden of both AKI and CKD, and to understanding pathways that drive progressive kidney fibrosis.

Fibrosis is the principal pathological process that underlies all progressive renal disease. It results from an imbalance between extracellular matrix (ECM) deposition and degradation leading to excessive fibrous scar accumulation and renal parenchymal damage. α-smooth muscle actin positive (α-SMA^+^) myofibroblasts are the principal effector cells that drive fibrosis and are derived from differentiation of stromal cells including resident fibroblasts and pericytes under the influence of pro-fibrotic cytokines, such as Transforming Growth Factor-β1 (TGF-β1)^10–15^. Fibroblasts have a common stromal progenitor cell-lineage but display considerable phenotypic plasticity. In fact, recent advances in lineage-tracing and single-cell RNA sequencing have confirmed the existence of distinct sub-populations of renal fibroblasts that are transcriptionally, functionally, and spatially heterogeneous^16–18^. However, it has been difficult to sufficiently enrich and isolate these subpopulations from kidneys to undertake studies to fully understand their biology. These stromal cells are key players in maintaining homeostasis and in facilitating repair and regeneration. In renal development they are critical for differentiation and maturation of nephrons. In the regenerative field, stromal cells assembled in co-culture with nephron and ureteric bud progenitors leads to the formation of organoids that closely resemble organotypic renal architecture and function, including the formation of nephron tubules^19^. In contrast, maladaptive fibroblast responses drive fibrosis and progressive CKD. Despite the increasing worldwide prevalence of renal disease there is a limited understanding of fibroblast heterogeneity in the kidney and the factors that regulate phenotypic variations. Therefore, insight into the distinct differences in fibroblast subpopulations during embryonic development and health; and how these can be recapitulated to promote renal recovery following injury is crucial to further advances in this field.

Hyaluronan (HA) is an extracellular matrix glycosaminoglycan that is critical in renal development and demonstrates increased expression in the renal stroma during progressive renal disease^20–22^. Our previous cell studies indicate that distinct alterations in pericellular HA generation, assembly and interactions are key to determining fibroblast heterogeneity^23–27^. HA is synthesized at the cell membrane by three HA Synthase (HAS) isoenzymes termed HAS1, HAS2 and HAS3. These isoenzymes are located on distinct chromosomes and are conserved in evolution suggesting that they may have unique and important roles in biology. In this study we show that HAS1 and HAS2 drive distinct fibroblast phenotypes *in vitro* and are identified in distinct subpopulations of interstitial stromal cells *in vivo* linked with fibrosis progression versus resolution following IRI. The results provide novel insights into the HAS isoenzymes as distinct regulators of stromal responses in progressive renal injury.

## METHODS

### Animals

Experiments were performed in line with institutional and UK Home Office guidelines under the authority of an appropriate project license. Adult male Lewis rats (8 to 12 weeks) were supplied by Charles River (Cambridgeshire, UK). All experimental procedures were carried out in accordance with local policies at JBIOS, Cardiff University and licensed under the Animal (Scientific Procedures) Act 1986, issued by the UK Government Home Office Department. Rats were housed in conventional housing with readily available chow and water. One day (24 h) prior to surgery, Buprenorphine (Temgesic) was added to drinking water to provide analgesia. Isoflurane anaesthesia was delivered via an induction chamber. The ARRIVE guidelines were followed for all *in vivo* experiments. Based on previous work, a power calculation indicated n=8 per group to give adequate power (alpha 0.05, beta 0.8).

#### IRI model

The central abdomen was shaved and a midline laparotomy incision made. Bilateral IRI was performed at 12 weeks of age by clamping both renal pedicles for 45 minutes and kidneys visually assessed for signs of ischaemia. The clamps were then removed, wound closure performed and animals recovered^28^. Kidney tissues were retrieved under terminal anaesthesia at 2 days (48 h), 7 days, 14 days and 21 days post-operatively. Blood samples for serum creatinine were taken at the same times. Kidneys were divided longitudinally with one half stored in formalin for histological evaluation and one half placed in RNALater (Thermofisher Scientific). Sham animals underwent a midline laparotomy alone with no renal pedicle clamping. Kidney tissues were retrieved, and blood samples undertaken under terminal anaesthesia at 48 hours, 7, 14 and 21 days post-operatively. A minimum of 8 rats were used per group according to appropriate power calculations. Schematic demonstrated in Figure 1A.

**Figure 1.**
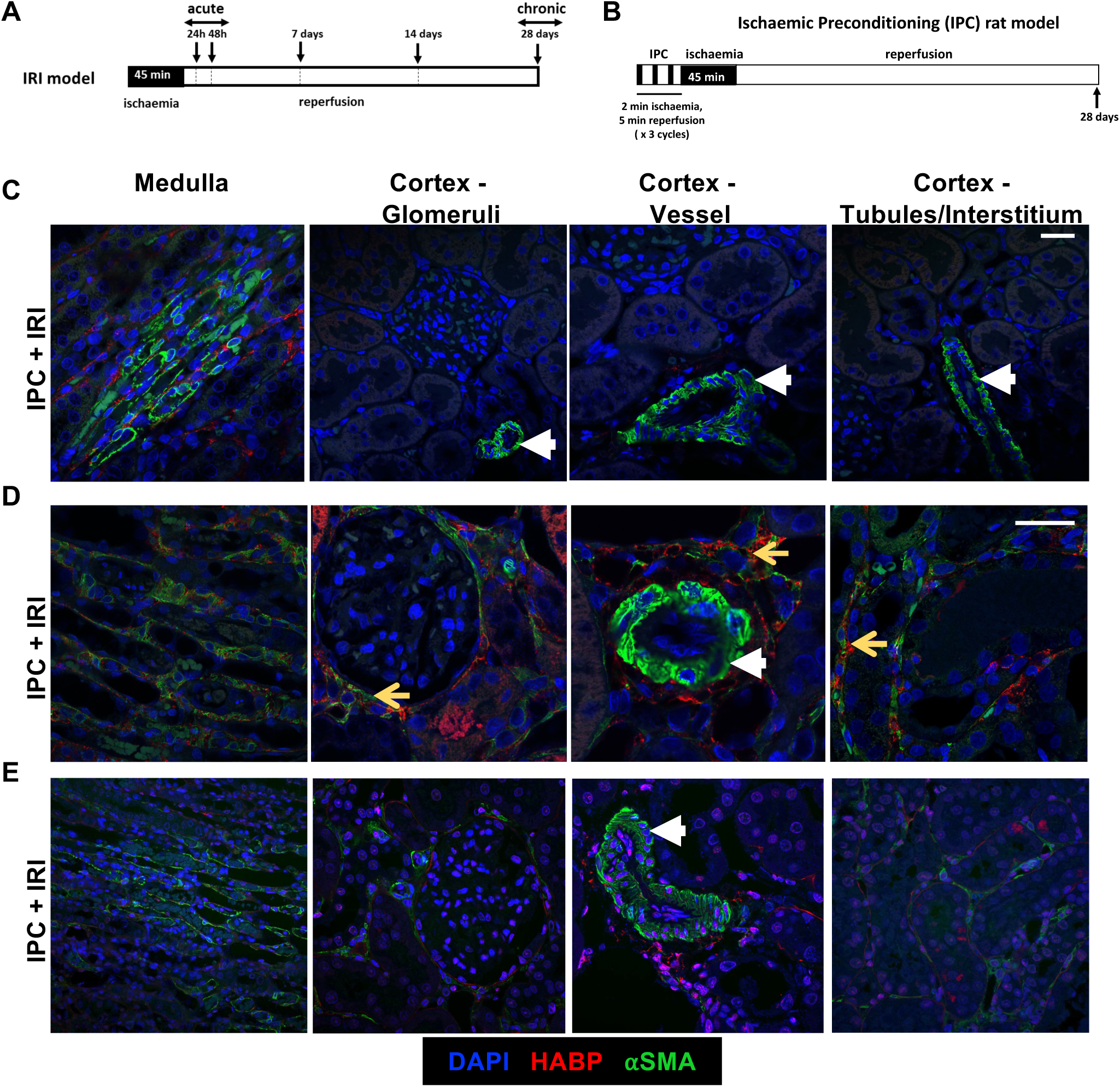
Increased HA deposition in the renal interstitial stroma following progressive damage correlates with increased myofibroblast positive regions. Schematic of (A) Ischaemia-Reperfusion Injury (IRI) model in ra and (B) Ischaemic Pre-Conditioning (IPC) model in rats. Adult male Lewis rats (n=72) underwent a midline laparotomy and were divided into two groups; sham operation (n=35), bilateral IRI (cross-clamping of both renal pedicles for 45 minutes) (n=37) or IPC+IRI (3 cycles of ischaemic pre-conditioning prior to IRI: 2 mins ischaemia and 5 mins reperfusion) (n=24). The groups were further subdivided depending on their post-operative observation time: 24h (sham n=8, IRI n=8, IPC+IRI n=8), 48h (sham n=11, IRI n=13), 7 days (sham n=4, IRI n=4, IPC+IRI n=4), 14 days (sham n=4, IRI n=4, IPC+IRI n=4) and 28 days (sham n=8, IRI n=8, IPC+IRI n=8). At the designated post-operative time, kidney tissue was recovered under terminal anaesthesia. (C) Kidney immunofluorescence sections in (C) normal (sham), (D) IRI and (E) IPC+IRI demonstrating HA (red), αSMA (green) and nuclear (blue) staining. White Arrowheads denote αSMA (green) staining of vessel walls. Yellow arrows denote areas of combined αSMA (green) and HABP (red) staining in the renal interstitium. Scale bars 20 μm, Original Magnification x400.

#### IPC model

A group of rats underwent ischaemic pre-conditioning (IPC) prior to IRI. In this group bilateral renal pedicles were clamped for 2 minutes (ischaemia), followed by release of clamps for 5 minutes (reperfusion)^28^. This process was performed three times in total, prior to bilateral IRI (cross clamping of both renal pedicles for 45 minutes). A group of animals underwent IPC alone with no subsequent IRI to investigate the effects of IPC alone. Kidneys tissues were retrieved, and blood samples undertaken as above at 2, 7, 14 and 21 days post-operatively and placed in RNALater or formalin. A minimum of 8 rats were used per group according to appropriate power calculations. Schematic demonstrated in Figure 1B.

#### Has1/3^-/-^ and wild-type Mice

Mouse kidneys from wild-type (WT) and *Has1/3* double knockout (KO) C57BL/6 mice were supplied by Professor Anna Plaas (Rush University, Chicago) and approved according to Rush University IACUC. Breeding, experimental procedures and maintenance were undertaken as previously described^29^. Mice had free access to water and chow and were exposed to the same light cycles (12 h light, 12 h dark) during the growth period. Mice were sacrificed at 12 weeks of age by CO_2_ asphyxiation and cervical dislocation. Kidneys were harvested and placed in formalin or RNALater for histological or gene expression analysis.

#### Embryonic Studies

Embryos were collected with the day of the vaginal plug designated as embryonic (E) day 0.5 and staged by morphologic criteria, which included somite number and eye and limb morphology. Kidneys were dissected from embryonic (E) day 15.5 embryos, a time at which all progenitor populations and maturing cell types co-exist, using a stereomicroscope.

### Primary human cells

#### Scarring fibroblast (SF) and non-scarring fibroblast (NSF) phenotypes

Donor-matched samples of dermal (scarring) and oral mucosal (non-scarring) fibroblasts were obtained by biopsy from consenting adults undergoing routine minor surgery (ethical approval obtained from South-East Wales Research Ethics Committee). The cells were isolated as previously described^26^.

#### Cell Culture

The above cells were cultured in Dulbecco’s modified Eagle’s medium and F-12 medium containing 2 mM L-glutamine, 100 units/ml penicillin, and 100 μg/ml streptomycin supplemented with 10% fetal bovine serum (FBS) (Biologic Industries Ltd., Cumbernauld, UK). The cells were maintained at 37 °C in a humidified incubator in an atmosphere of 5% CO_2_, and fresh growth medium was added to the cells every 3– 4 days until confluent. The cells were incubated in serum-free medium for 48 hours before use in experiments, and all experiments were done under serum-free conditions unless otherwise stated. All experiments were undertaken using cells at passage 6 –10. Experimental conditions were designed such that cells were either incubated in serum-free medium (control) or in medium containing 10 ng/ml TGF-β1 for 72 hours.

#### Construction of Adenoviral Vectors and NSF Infection

Adenoviral vectors for enforced expression of HAS1 and HAS2 open-reading frames (ORFs) were constructed by PCR amplification (Table 1 Supplementary data) of the HAS ORFs from the plasmid expression vectors described above, followed by homologous recombination into the replication deficient, E1/E3 deleted, adenoviral vector pAdZ-CV5 as described elsewhere^30^. Primary OMFs were grown to 70% confluence and growth arrested for 24 h in serum free medium. Viral vectors were added to the cells in a minimal volume of serum free medium and rocked for 2 hours before replacement with serum free medium for 48 hours and subsequent experimentation with serum free medium alone or containing 10 ng/ml TGF-β1.

#### siRNA Knockdown of Gene Expression

Annealed oligonucleotide siRNA reagents for the knockdown of HAS1 (assay ID 119443), HAS2 (s6458), and scrambled negative control transfection (4611) were purchased from Life Technologies and used in accordance with the manufacturer’s guidelines. A final concentration of 30 nM of each siRNA was transfected into cultured cells using Lipofectamine 2000 (Life Technologies), as described previously^23^.

### Reverse transcription (RT) and quantitative PCR (qPCR)

qPCR was used to assess mRNA expression levels of *HAS1, HAS2*, *Cdh1* (E-Cadherin), *ACTA2 (*α-SMA) (), *HYAL2 (*hyaluronidase 2), *FN1 (*Fibronectin), *EDA-FN* (extra domain-A–fibronectin), *COL1A1 (*collagen type I alpha 1 chain) and *FAP* (Fibroblast Activated Protein)). Total RNA was extracted from frozen kidney tissue stored in RNALater with RNeasy mini kit (Qiagen), quantified by NanoDrop spectrophotometer and used to prepare cDNA with Multiscribe reverse transcriptase (Thermo Fisher Scientific). Quantitative PCR was performed using Power SYBR Green or Taqman PCR Master Mix (Thermo Fisher Scientific) and specific primers (supplemental Table 1). Samples were analyzed on a ViiA 7 Real-Time qPCR System (Thermo Fisher Scientific), and relative expression levels were normalized to a standard reference gene (18s rRNA) using the 2^−ΔΔCT^ method^23^.

### Histology, Immunohistochemistry (IHC) and Immunofluorescence (IF)

Fixed rat or mouse kidneys were processed for embedding in paraffin and cut into 5 *μ*m sections for Masson’s Trichrome staining, HA binding protein (HABP) staining or immunostaining. Deparaffinized kidney sections were rehydrated in graded alcohols (100%, 96%, 70%, and 50%), and antigen retrieval was performed in sodium citrate buffer in the autoclave at 120°C for 20 minutes. For IHC signal detection, endogenous peroxidase activity was quenched by incubation in 3% (vol/vol) H_2_O_2_ /methanol for 10 minutes at room temperature. Where biotinylated secondary antibodies were used, sections were incubated with avidin/Biotin block (Vector Laboratories) for 10 minutes. Sections were blocked in 10% goat serum (Vector Laboratories) for 20 minutes at room temperature and incubated overnight at 4°C with primary antibodies (Table 2 Supplementary Data). Immobilized antibodies were detected by UltraVision-LP HRP Polymer detection system specific for anti-mouse and anti-rabbit IgG (ThermoFisher Scientific), or with biotinylated secondary antibodies and AB reagents (Vector Laboratories) according to manufacturer’s instructions. DAB (Vector Laboratories) and hematoxylin were used as the chromogen and the nuclear counterstain, respectively. For detection of HA, a HA binding protein (HABP, Merck, cat.number 385911) with a biotin conjugate was used (1:200) followed by AB reagent or streptavidin conjugated with AlexaFlour 488/594 fluorophore (ThermoFisher Scientific). For IF detection, AlexaFluor 488 goat anti-mouse IgG (H+L) and Alexa Fluor 594 goat anti-mouse IgG (H+L) secondary antibodies were used at 1:500 dilution (ThermoFisher Scientific). DAPI (4’,6-diamidino-2-phenylindole) was used to visualize the nuclei. In negative controls, primary antibodies or HABP were omitted. Brightfield kidney sections were examined using a Leica DMLA Light Microscope equipped with an Olympus DP27 Microscope digital camera. For IF kidney sections were examined and imaged using a Zeiss LSM880 Airyscan Confocal Laser Scanning Microscope.

### Visualisation of Pericellular HA by Particle Exclusion Assay

Pericellular HA coats were visualized by the exclusion of horse erythrocytes as reported previously^26^. Formalized horse erythrocytes were washed in PBS and centrifuged at 1,000 × *g* for 7 min at 4 °C. The pellet was resuspended in serum-free medium at an approximate density 1 × 10^8^ erythrocytes/ml. A total volume of 500 μl of this suspension was added to each 35-mm dish containing subconfluent cells and swirled gently for even distribution. The dishes were incubated at 37 °C for 15 min to allow the erythrocytes to settle around the cells. Control cells were incubated with 200 μg/ml bovine testicular hyaluronidase in serum-free medium for 30 min prior to the addition of formalized horse erythrocytes. On settling, the erythrocytes were excluded from zones around the cells with HA pericellular coats. This was viewed under the microscope as an area of erythrocyte exclusion. Zones of exclusion were visualized on a Zeiss Axiovert 135 inverted microscope.

### Collagen gel contractility Assay

Type I collagen was extracted from rat-tail tendon as previously described^31^. Approximately 2.5 × 10^5^ fibroblasts/mL were mixed into collagen lattice–forming solutions (2.5 mL 20% v/v fetal calf serum– Dulbecco’s modified Eagle’s low-glucose medium, 500 μL of 0.1 mol/L NaOH, and 2 mg/mL type I collagen; total volume of 5 mL). Fibroblast-populated collagen lattices (FPCLs) were maintained at 37°C, in a 5% CO_2_ atmosphere for 1 hour, for collagen polymerization to occur. FPCLs were gently detached from the plate edges and resuspended in serum-free medium containing appropriate treatments. FPCLs were measured at 0, 24, 48, 72, 96, 120, 144 hours after initial lattice fabrication. The mean FPCL contraction values were obtained from analysis by ImageJ software version 1.37c.

### Scratch-Wound Migration Assay

Scratching quiescent fibroblasts cell monolayers with sterile 200-μL pipette tips generated linear denuded areas. The cells were gently washed with PBS to remove detached cells and then replenished with fresh serum-free medium containing appropriate treatments. The wound width was photographed at 0, 24, 48, and and 72 hours or until closure, using an Axiovert 100 mol/L inverted microscope (Carl Zeiss, Oberkochen, Germany) fitted with a digital camera (Orca-1394; Hamamatsu Photonics K.K., Hamamatsu, Japan). Measurements were obtained using ImageJ software version 1.37c and the area of wound closure was measured to assess the rate of migration.

### *In silico* sequence analysis

A region totalling 3 kb upstream of the HAS isoforms’ transcription start sites was extracted from the Ensembl genome browser (http://www.ensembl.org), taking sequence from position −1 to −3,000 with respect to the transcription start sites, and were analysed for conserved noncoding regions (CNRs) using VISTA^32^.

### Statistical Analysis

The two-tailed, unpaired *t*-test was used to assess statistical differences between the two experimental groups. For experiments with multiple experimental groups, one-way analysis of variance was used to identify statistical differences across groups, followed by post-test Bartlett and multiple comparisons. For experiments with multiple experimental conditions, two-way analysis of variance was used, followed by post-test Tukey multiple comparisons. Graphical data are expressed as means ± standard error of mean (SEM). All data were analyzed using GraphPad Prism software version 6 (GraphPad Software, San Diego, CA). *P* < 0.05 was considered statistically significant.

## RESULTS

### HA demonstrates increased stromal deposition following IRI and localises in areas of α-SMA^+^ myofibroblast expansion

Our studies demonstrated that IRI (Figure-1A) led to both acute renal injury and chronic interstitial fibrosis. In contrast, we showed that ischaemic preconditioning (IPC) prior to IRI (Figure-1B) led to enhanced recovery from IRI acutely and from renal interstitial fibrosis chronically^28^. To determine the renal expression and localisation of HA, IF was performed in rat kidney sections 48h following IRI. HA has low immunogenicity and cannot be detected directly *via* an antibody but is detectable through incubation with a specific HA Binding Protein (HABP) derived from bovine nasal cartilage. Our previous *in vitro* data in fibroblasts indicated that the generation of pericellular HA matrices is critical for differentiation-to and maintenance-of pro-fibrotic α-SMA^+^ myofibroblast phenotypes^26^. Hence co-staining for α-SMA was performed to determine the relationship between the localisation of HA and α-SMA^+^ myofibroblasts *in vivo*.

The results demonstrated that in healthy (sham) kidneys there was HA (red) present in the renal medulla surrounding the loops of Henle. Previous studies indicate that medullary HA may be involved in urine concentration^33^. Minimal HA was evident in the renal cortex either in the glomerular, perivascular or tubular/interstitial areas. Furthermore, there was no interstitial α-SMA^+^ (green) staining in healthy kidneys indicating absence of pro-fibrotic myofibroblasts. The α-SMA staining in healthy kidneys was limited to the walls of the renal arteries/arterioles, where it stained contractile α-SMA^+^ vascular smooth muscle cells (Figure-1C). Following IRI, HA deposition was increased in the renal cortex and was particularly evident in peri-glomerular, perivascular and peri-tubular interstitium. Staining for α-SMA^+^ myofibroblasts was also abundant in these HA-enriched areas (Figure-1D), supporting our previous data demonstrating HA as essential for differentiation to a myofibroblast phenotype. Application of IPC prior to IRI resulted in a considerable change to HA and α-SMA localisation in the renal cortex, with only modest expression of both now present within the renal stroma (Figure-1E).

### HAS1 localises in areas of abundant HA generation in the stroma of developing mouse kidney

HA has an established role in embryonic development, with HAS2^-/-^ mice demonstrating lethal cardiovascular anomalies^34^. To gain insights into the role of HA in kidney development, E15.5 mouse kidneys were stained for HA. At E15.5 the overall nephron segments are specified, but new nephrons continue to be generated from nephron progenitors^35^. Therefore, insights into the stromal environment that supports nephron regeneration was assessed at this stage to enable subsequent comparisons to be made with stroma in mature kidneys. The results demonstrated abundance of HA throughout the developing kidney with expression evident in the nephrogenic zone (nz), surrounding the ureteric bud (ub) branches, in the cortical stroma (cs) grouped around developing tubules, and in the medullary stroma (ms) (Figure-2A).

**Figure 2.**
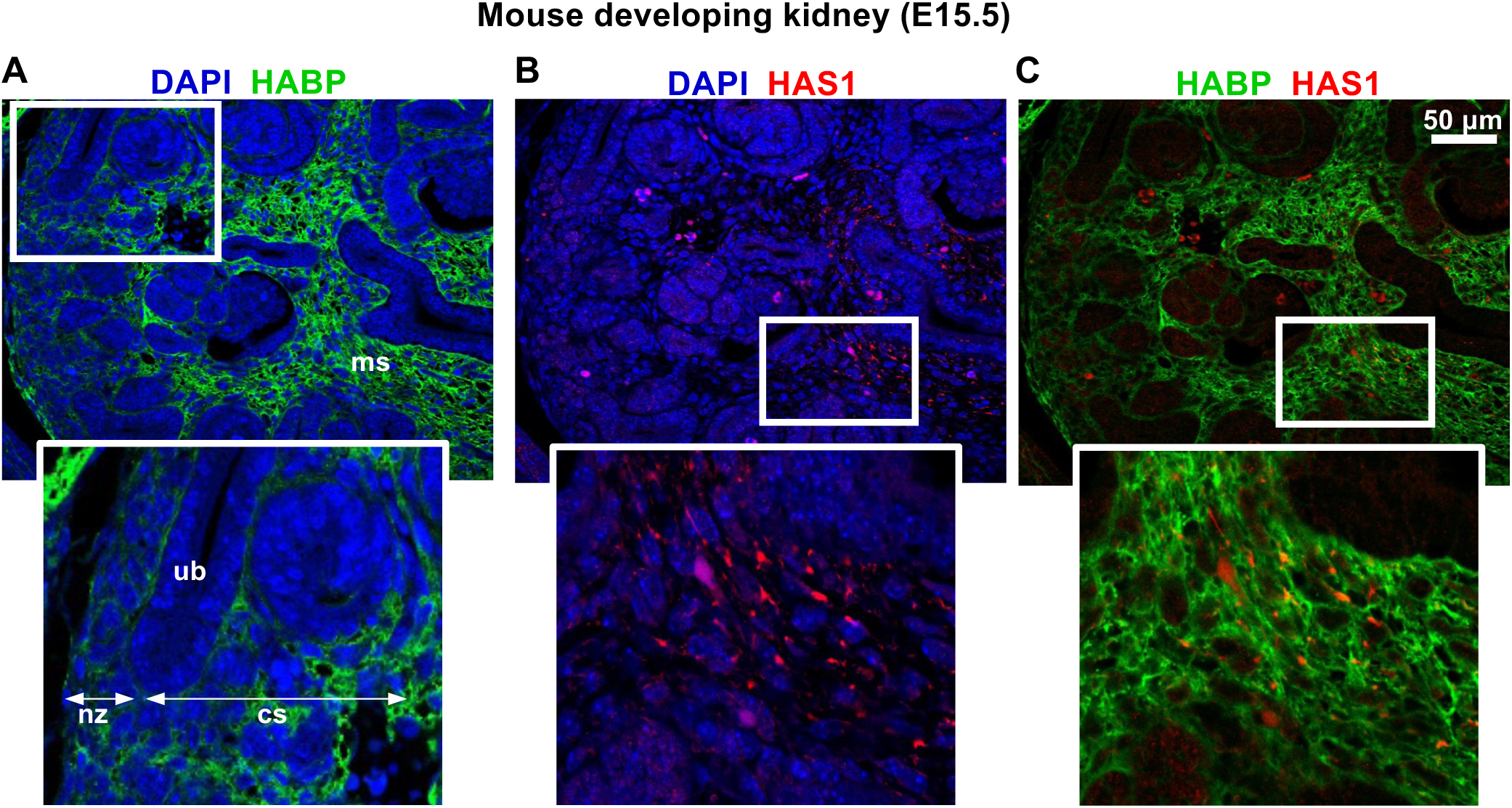
Localisation of HA in developing kidneys. Immunofluorescence detection of (A) hyaluronan (HA binding protein, HABP) (green) and (B) HAS1 (red) in mouse foetal kidneys demonstrating abundance of HA and HAS1 in the stroma of the nephrogenic zone (nz) surrounding ureteric bud (ub) branches and induced nephrons in the cortical stroma (cs) grouping around sets of tubules, and in the stromal interstitium of the medulla in mouse E15.5 kidneys. (C) Co-localisation of HAS1 (red) and HA (green). White boxed regions exhibit sections with higher magnification. Scale bars 50 μm, Original Magnification x400.

To determine the HAS isoenzyme responsible for HA generation in kidney development, HAS1 and HAS2 IHC was performed. HAS1 was detected in a subset of tubular cells (supplementary data Figure-1A). In contrast, detection of HAS2 was negative (supplementary data Figure-1B). IF analysis demonstrated co-localisation of HA (green) with HAS1 (red) in the stroma (Figure-2B&2C) suggesting HAS1 plays a prominent role in stromal HA generation in kidney development.

### The spatiotemporal distribution of HA in injured mature kidneys resembles that seen during embryonic kidney development

Staining for HABP was undertaken in healthy kidneys and in kidneys following 24h to 28d post-IRI. Our recent studies indicate that in this model there is prominent acute renal injury at 24h and 48h, which begins to subside at 7d. At 24h, 48h and 7d there is also evidence of tubules undergoing repair; whilst at 14d post-IRI there is limited evidence of acute/chronic injury. At 28d post-IRI there is evidence of florid fibrosis and damage in the tubulointerstitium compared to sham kidneys^28^. In view of this we denoted the 24h and 7d period as the regenerative phase and the 28d period as the chronic fibrosis phase post-IRI. HA was absent from the renal cortex in health but was rapidly expressed in the renal stroma following ischaemia, with staining evident at 24h post-injury. Stromal HA was increased from 24h up to 7d and was prominent in areas surrounding repairing tubules and in capillary dense regions (black arrows). This strikingly resembled stromal HA distribution in the developing kidney (Figure-2), where stromal HA also surrounded areas of tubular development and new capillary formation. At 14d post-IRI, stromal HA was present but less prominent and by 28d, HA was only present in stromal matrices in areas of unresolved inflammation and fibrotic scarring (Figure-3).

**Figure 3.**
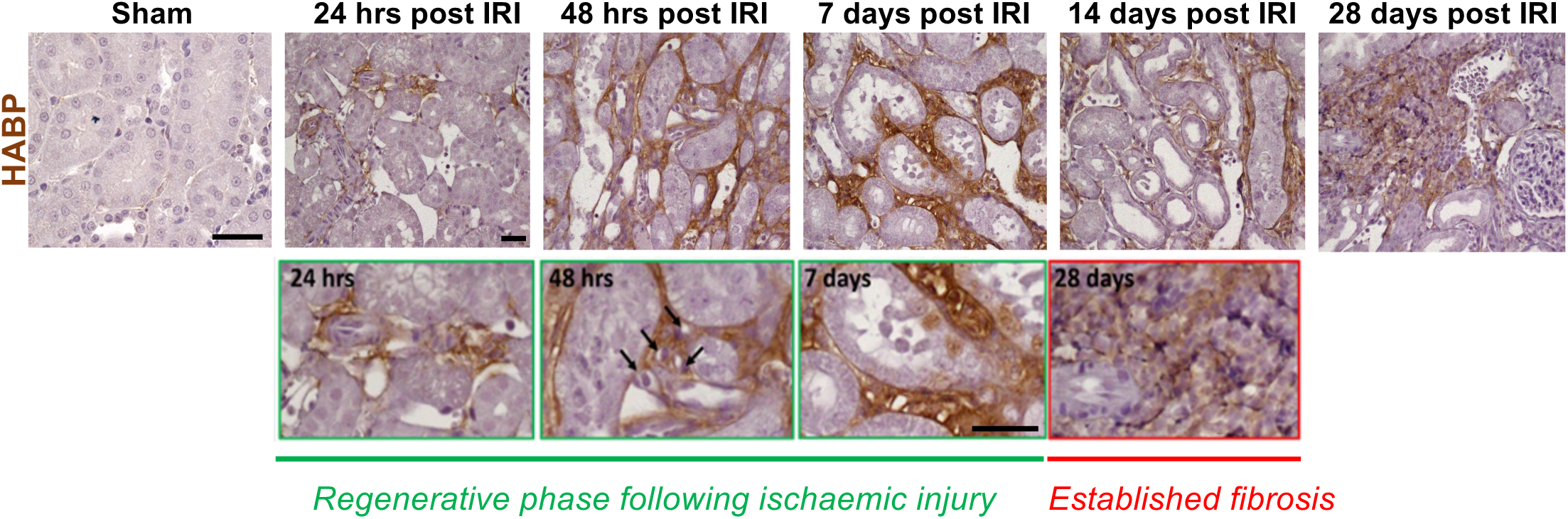
The spatiotemporal localisation and pattern of HA staining in mature rat kidneys. Histological staining of kidney sections with HA binding protein (HABP) from IRI rat model at 24 hours, 48 hours, 7 days, 14 days and 28 days post IRI. Scale bars 20 μm, Original Magnification x400.

### Distinct HA Synthase isoforms demonstrate differential patterns of expression in mature kidneys

Synthesis of HA is dependent on three membrane-bound HA synthase enzymes: HAS1, HAS2 and HAS3. The role of these distinct isoenzymes in the generation of stromal HA matrices in mature kidneys in health, following ischaemic injury and during recovery from ischaemic injury was investigated using the models of IRI +/- IPC described. Initially, mRNA expression of the HAS isoenzymes was assessed using RT-qPCR of whole kidney. The data demonstrated that there was 1.5-2-fold induction of *Has1* expression following IRI. In IPC+IRI animals, renal expression of *Has1* remained elevated. In contrast, *Has2* mRNA expression was markedly increased compared to *Has1* expression following IRI, with a 9-10-fold induction in expression. In IPC+IRI animals, *Has2* mRNA expression was attenuated, suggesting that HAS2 is more likely to be disease promoting in the kidneys than HAS1 (Figure-4A). *Has3* mRNA expression was unchanged across all cohorts (data not shown).

**Figure 4.**
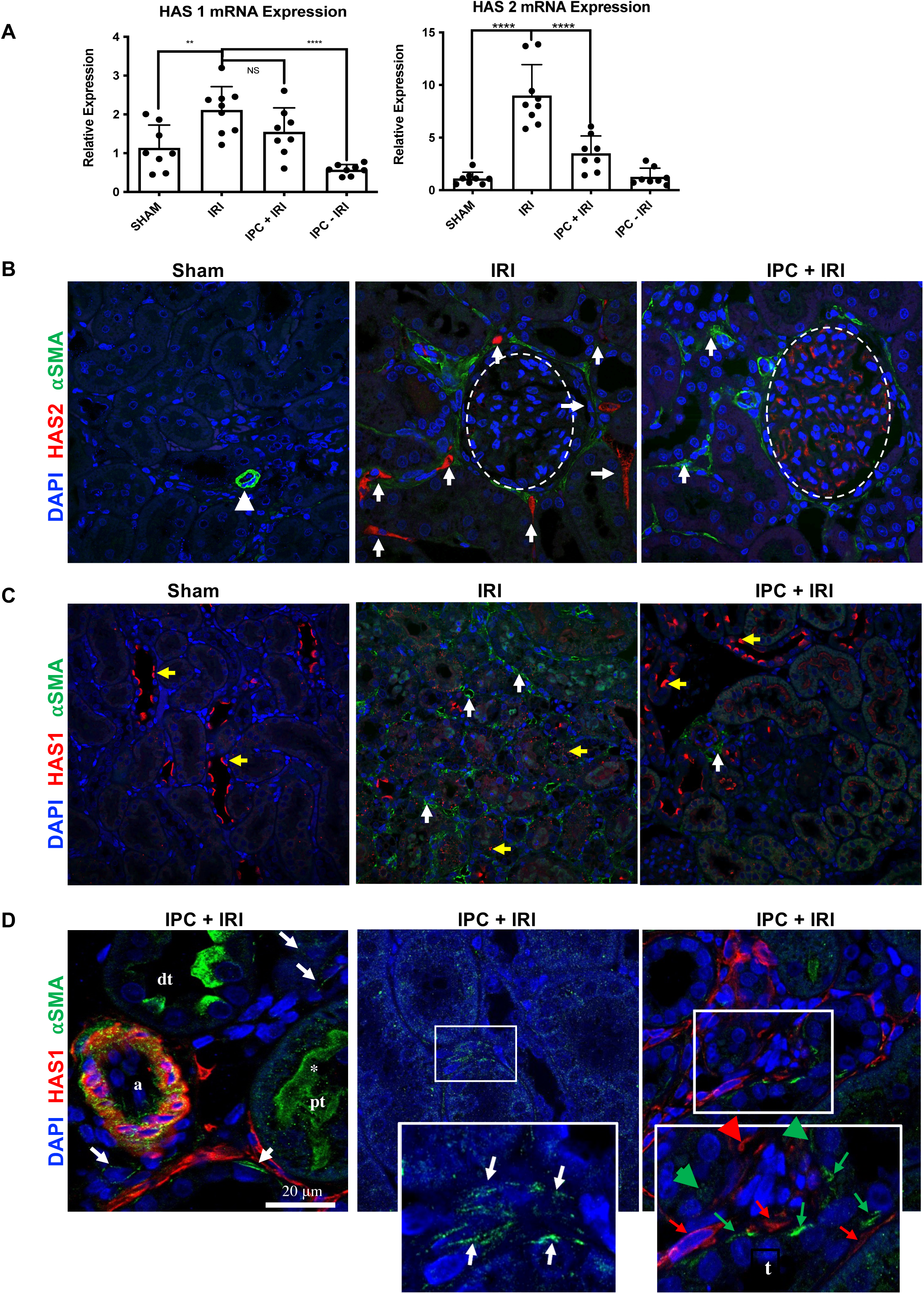
HAS1 and HAS2 demonstrate differential expression patterns in rat kidneys. (A) Kidneys from normal (Sham), IRI alone and IPC+IRI models were assessed using RT-qPCR of whole kidney to assess expression of *HAS1* and *HAS2* mRNA. (B) Immunofluorescence detection of HAS2 (red) and α-SMA (green) in normal (sham), IRI only and IRI pre-treated with IPC rat kidneys. Nuclei are stained with DAPI (blue). White arrowhead denotes staining of small blood vessels with α-SMA. White arrows denote HAS2 staining of stromal cells in the interstitium. Dashed white circles highlight the glomeruli. (C) Immunofluorescence detection of HAS1 (red) and α-SMA (green) in normal (sham), IRI only and IRI pre-treated with IPC rat kidneys. Nuclei are stained with DAPI (blue). White arrows denote increased α-SMA (green) staining in areas of damage/fibrosis. Yellow arrows demonstrate HAS1 (red) expression at the apical aspect of a proportion of tubular epithelial cells. (D) Higher Magnification images of rat kidneys treated with IPC prior to IRI stained for HAS1 (green), α-SMA (red) and DAPI (nuclei, blue). Increased HAS1 positive staining can be identified in the arterioles (a) at the apical border of distal tubular epithelial cells (dt) and in interstitial stromal cells (white arrows). In the last panel α-SMA positive stromal cells (red) and HAS1 positive stromal cells (green) are identified by red and green arrowheads respectively to distinguish them. Scale bars 20 μm, Original Magnification x400.

IF detection of HAS1 and HAS2 isoenzymes was subsequently undertaken. Initially, dual-staining compared HAS2 (red) and α-SMA^+^ (green) myofibroblast localisation (Figure-4B). As previously demonstrated, α-SMA^+^ staining was observed only in the vessel walls in sham animals. However, following IRI, α-SMA^+^ staining was identified throughout the interstitium. In IPC+IRI animals, interstitial α-SMA^+^ staining was attenuated. In sham animals, HAS2 was not identified in the renal cortex, whilst there was a noticeable increase in interstitial HAS2 in IRI animals. The HAS2^+^ cells localised to regions enriched with α-SMA^+^ cells. In accordance with Has2 gene expression, HAS2 protein expression was also attenuated in the interstitial stroma in IPC+IRI animals.

HAS1 (red) localisation in the kidney were explored next (Figure-4C). In sham kidneys HAS1 was only present at the apical border of tubular epithelial cells in a subset of tubules comprising approximately 30-40 percent of the renal cortex. Following IRI, HAS1 immunoreactivity was decreased, and the distinct localisation seen in the sham animals was disrupted. As expected, extensive interstitial α-SMA (green) was also present following IRI; however, unlike with HAS2, α-SMA and HAS1 were not identified in the same areas, with no HAS1 demonstrated in the interstitial space. In animals pre-treated with IPC, HAS1 demonstrated apical tubular cell expression in a subset of tubules and this was marginally enhanced compared to sham/healthy kidneys. Moreover, HAS1 was also present in other non-tubular cell-types including expression in vascular smooth muscle cells and in a subpopulation of interstitial stromal cells comparable to what was observed during kidney development. Of note, interstitial HAS1^+^ cell staining in IPC+IRI treated animals was distinct from α-SMA^+^ interstitial cell staining (red *versus* green arrows) (Figure-4D).

### HAS1^+^ and HAS2^+^ fibroblasts display distinct pericellular matrices

Our subsequent aim focussed on understanding cellular functions of HAS1 and HAS2. Compared to other adult tissues in the human, the oral mucosa undergoes scarless healing^36^. Furthermore, fibroblasts from the oral mucosa demonstrate intrinsic biological properties that explains their propensity for scar-free healing. Specifically, oral mucosal fibroblasts demonstrate resistance to TGF-β1-driven myofibroblast differentiation and proliferation^26, 37^. Hence, to better understand the cellular biology involved in scarring/fibrosis *versus* scar-free healing, we used donor-matched oral mucosal and dermal fibroblasts as models of non-scarring and scarring fibroblast phenotypes respectively. We also previously showed that the TGF-β1 driven responses in oral mucosal and dermal fibroblasts are causally-linked to the cells ability to generate pericellular HA matrices, which have been established as being essential for differentiation-to and maintenance-of the pro-fibrotic α-SMA^+^ myofibroblast phenotype^24, 26, 37^. Through these cumulative studies we fully characterised the HA profile of these distinct fibroblast phenotypes and showed that dermal fibroblasts are able to express both HAS1 and HAS2 isoenzymes, whereas oral fibroblasts do not express HAS1 and have low-level HAS2 expression^26^. In this study, we used adenoviral vectors to over-express either HAS1 or HAS2 in oral mucosal fibroblasts and thereby evaluate the effect of manipulating these distinct iso-enzymes on TGF-β1 responses in fibroblasts. We also used siRNA to abrogate HAS1 and/or HAS2 in dermal fibroblasts. Initial experiments were undertaken to confirm HAS1 and HAS2 over-expression and/or knockdown (supplementary data Figure-2).

The red-cell exclusion assay was used to assess effects of forced HAS1 *versus* HAS2 over-expression in oral fibroblasts on the formation of HA pericellular matrices. HAS1^+^ fibroblasts generated small HA pericellular matrices, whereas HAS2^+^ fibroblasts demonstrated larger HA pericellular matrices (Figure-5A). Coat-size data were displayed for quantification (Figure-5B). RT-qPCR was used to assess gene expression markers of other non-HA matrix components in HAS1^+^ *versus* HAS2^+^ cells. HAS2^+^ fibroblasts had significantly enhanced mRNA expression of EDA-fibronectin (*EDA-FN*) and Collagen-AI (*COL1A1*). In contrast, HAS1^+^ fibroblasts demonstrated higher expression of non-EDA fibronectins and but had lower levels of *EDA-FN and COL1A1* (Figure-5B). Conversely, siRNA-mediated knockdown of HAS1 *versus* HAS2 undertaken in dermal fibroblasts demonstrated that HAS2 knockdown led to marked attenuation of *EDA-FN* and *COL1A1*, whereas HAS1 knockdown led to marked attenuation of *FN1* expression (figure-5D).

**Figure 5.**
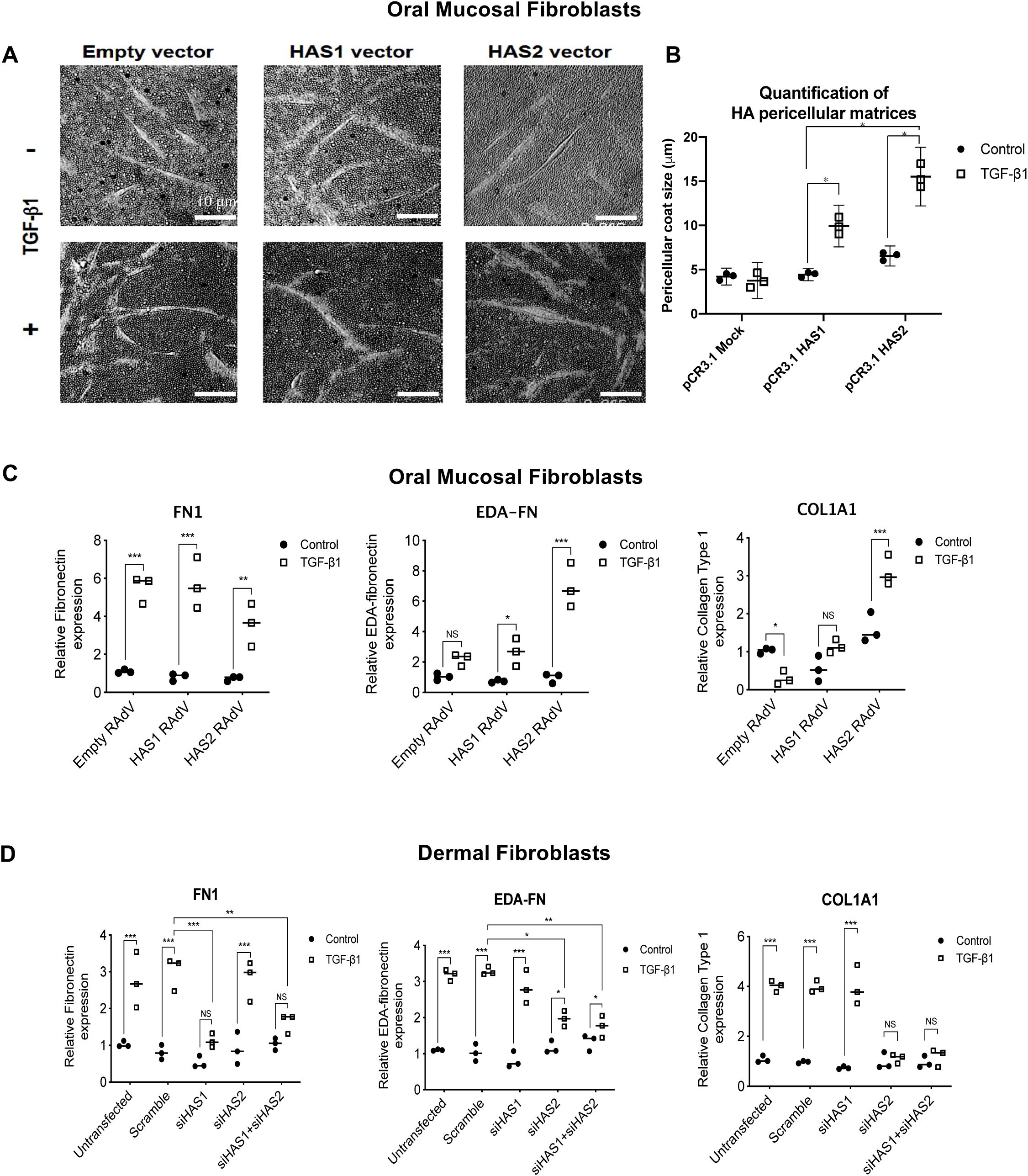
HAS1 and HAS2 isoenzymes are associated with distinct pericellular matrices in fibroblasts. (A) Red cell exclusion assay: The effect of 10 ng/ml TGF-β1 on pericellular HA coat formation in OMFs transfected with empty plasmid vector pCR3.1, or pCR3.1 with inserted HAS1 or HAS2 ORFs, was visualized using formalin-treated horse erythrocytes and observed by light microscopy. (B) Pericellular HA coat sizes were calculated by image analysis of OMFs following the HAS expression vector treatments described above in the absence (●) or presence (□) of 10 ng/ml TGF-β1. (C) Empty recombinant adenoviral vector (RAdV) and HAS1 or HAS2 RAdVs were used to infect oral mucosal fibroblasts in the absence or presence of 10 ng/ml TGF-β1 for 72 hours. RT-qPCR was used to assess mRNA expression of Fibronectin (*FN1*), EDA-splice variant fibronectin (*EDA-FN*) and collagen-1 (*COL1A1*). (D) Dermal fibroblasts were transfected with scrambled siRNA or siRNA targeting HAS1, HAS2 or both and incubated in the absence or presence of 10ng/ml TGF-β1 for 72 hours. RT-qPCR was used to assess mRNA expression of Fibronectin (*FN1*), EDA-splice variant fibronectin (*EDA-FN*) and collagen-1 (*COL1A1*). Data are presented as dot plots representative of 3 separate experiments. \**P*<0.05; \*\**P*<0.01; \*\*\**P*<0.001.

### HAS1^+^ and HAS2^+^ fibroblasts display distinct TGF-β1-driven phenotypes

Gene expression of two stromal markers were compared in HAS1^+^ *versus* HAS2^+^ fibroblasts. mRNA expression α-SMA (*ACTA2*) was assessed to determine differentiation to the classical pro-fibrotic myofibroblast phenotype. In addition, mRNA expression of the cell-surface protease Fibroblast Activated Protein (*FAP*) was assessed. FAP has been identified as a stromal marker that provides protection from fibrotic damage in lung fibrosis, through its ability to break down extracellular matrix^38^. In our cells, forced HAS1 over-expression in oral fibroblasts led to only a small increase in *ACTA2* expression, whereas forced HAS2 over-expression led to over a 4-fold increase in *ACTA2* expression (Figure-6A). In dermal fibroblasts, HAS1 knockdown had only a marginal effect on *ACTA2* expression, whereas HAS2 knockdown led to a marked attenuation of *ACTA2* expression (Figure-6B). Conversely, forced HAS1 over-expression enhanced *FAP* levels in oral fibroblasts compared to forced HAS2 expression (Figure-6C). Moreover, HAS1 knockdown had a more pronounced effect on *FAP* mRNA expression in dermal fibroblasts compared to HAS2 knockdown (Figure-6D). Interestingly, incubating scar-forming fibroblast phenotypes with TGF-β1, led to limited/no upregulation of *HAS1* mRNA. In contrast, stimulation with the anti-fibrotic growth factor BMP7 led to marked *HAS1* upregulation; indicating HAS1 as a downstream mediator of BMP7 that may be involved in its reno-protective role in the kidneys (Figure-6E).

**Figure 6.**
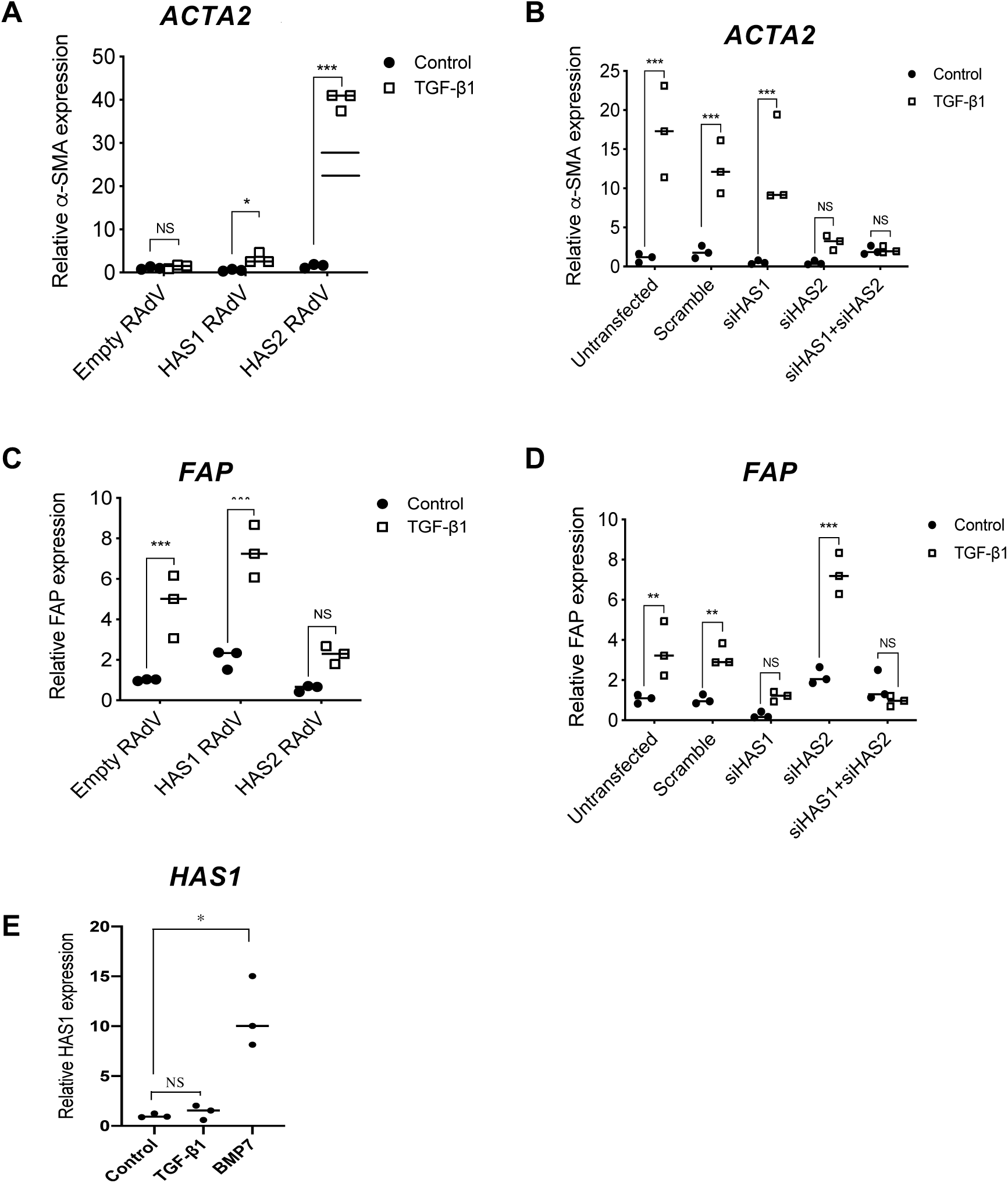
HAS1 and HAS2 isoenzymes in fibroblasts are associated with distinct expression patterns of α-SMA and FAP. (A) Empty recombinant adenoviral vector (RAdV) and HAS1 or HAS2 RAdVs were used to infect oral mucosal fibroblasts in the absence (●) or presence (□) of 10 ng/ml TGF-β1 for 72 hours. RT-qPCR was used to assess mRNA expression of α-SMA. (B) Dermal fibroblasts were transfected with scrambled siRNA or siRNA targeting HAS1, HAS2 or both and incubated in the absence or presence of 10ng/ml TGF-β1 for 72 hours. RT-qPCR was used to assess mRNA expression of α-SMA. (C) Empty recombinant adenoviral vector (RAdV) and HAS1 or HAS2 RAdVs were used to infect oral mucosal fibroblasts in the absence or presence of 10 ng/ml TGF-β1 for 72 hours. RT-qPCR was used to assess mRNA expression of *FAP*. (D) Dermal fibroblasts were transfected with scrambled siRNA or siRNA targeting HAS1, HAS2 or both and incubated in the absence or presence of 10ng/ml TGF-β1 for 72 hours. RT-qPCR was used to assess mRNA expression of *FAP*. (E) Dermal Fibroblasts are stimulated with either serum-free medium (control), 10ng/ml TGF-β1 or 200ng/ml of BMP7 for 72 hours. RT-qPCR is used to determine mRNA expression of HAS1. Data are presented as dot plots representative of 3 separate experiments. \**P*<0.05; \*\**P*<0.01; \*\*\**P*<0.001.

### HAS1^+^ fibroblasts are pro-migratory in comparison to HAS2^+^ fibroblasts that display a pro-contractile phenotype

A well-established characteristic of the pro-fibrotic α-SMA^+^ myofibroblast phenotype is its pro-contractile properties. Hence the role of HAS1 and HAS2 iso-enzymes on TGF-β1-driven contractility was investigated. Oral mucosal fibroblasts over-expressing HAS1 had attenuated contractility compared to oral fibroblasts expressing HAS2 (Figure-7A). To quantify this the diameter of collagen-gels was measured every 24h. The results indicated a statistically significant increase in collagen-gel contraction following HAS2 over-expression (Figure-7B).

**Figure 7.**
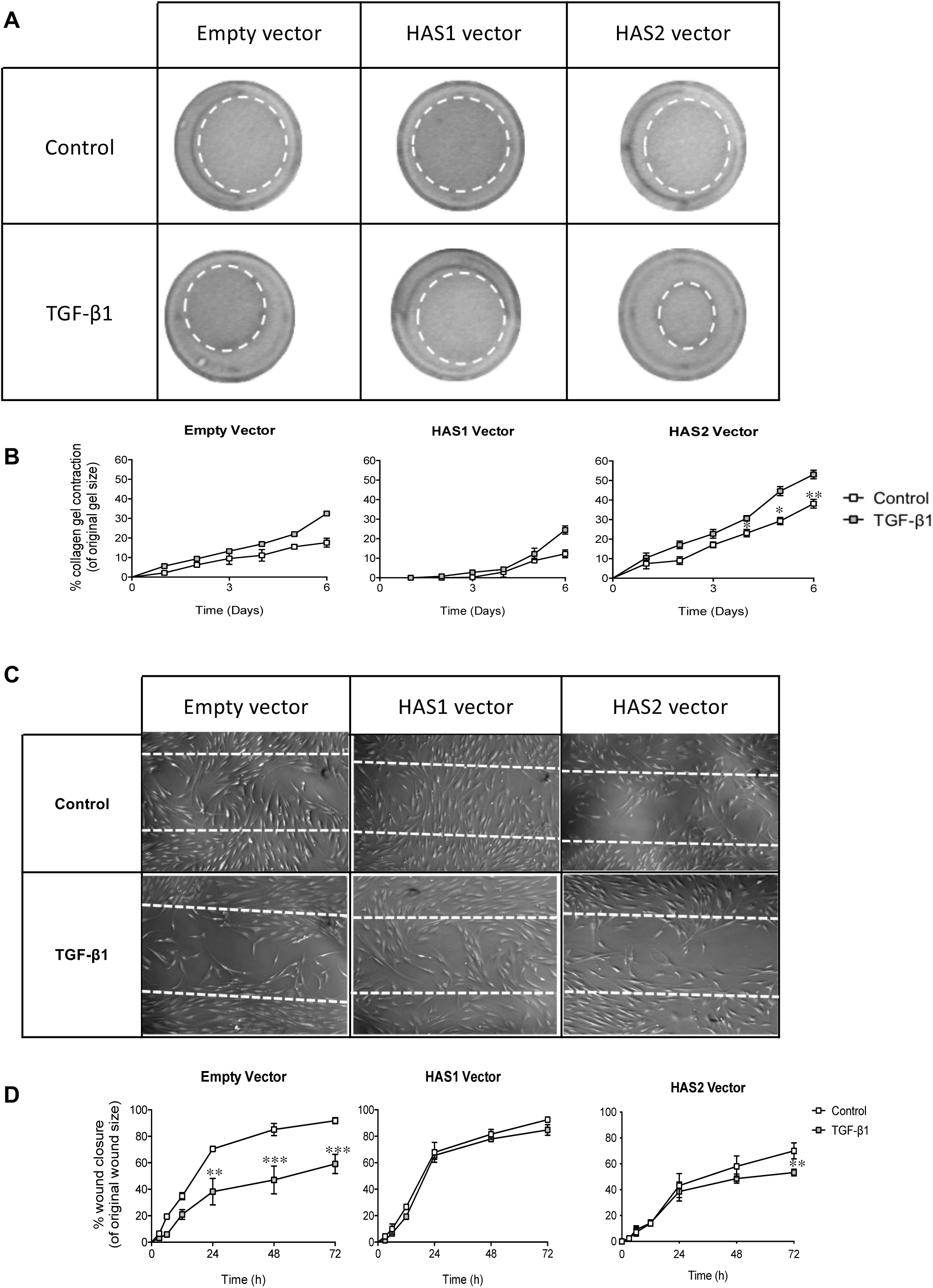
HAS1 and HAS2 isoenzymes have distinct effects on fibroblast contraction and migration. Empty recombinant adenoviral vector (RAdV) and HAS1 or HAS2 RAdVs were used to infect oral mucosal fibroblasts. (A) Cells were seeded into collagen gels and left to polymerize before incubation in the absence or presence of 10 ng/mL TGF-β1. Collagen gels were imaged over the annotated time points and measured for analysis of rate of contraction. Dotted circles indicate the measured gel areas at this time point (B) Percentage collagen gel contraction of original gel size is quantified at each time and annotated as a graph. (C) Cells were growth-arrested and scratched using a 20 μL pipette tip. After a phosphate-buffered saline wash, the cells were incubated in the absence or presence of 10 ng/mL TGF-β1. Scratch assays were imaged at the indicated time points and the area of closure was measured to assess the rate of migration. Dotted lines represent the original scratch edges. (D) Percentage wound closure compared to original wound size is measured to quantify rate of wound closure and annotated as a graph (D). Data are expressed as means ± SEM of three individual experiments. \**P*<0.05; \*\**P*<0.01; \*\*\**P*<0.001.

FAP facilitates stromal cell migration through matrix degradation in tissues^38^. In view of the data indicating enhanced *FAP* expression in Figure-6, the role of HAS isoenzymes fibroblast migration was assessed. The results demonstrated that TGF-β1 had an anti-migratory effect in un-transfected oral fibroblasts. HAS1 over-expression negated the TGF-β1 anti-migratory effects, whereas HAS2 overexpression had limited impact on this (Figures-7C &7D).

### Kidneys only expressing HAS2 are primed towards developing fibrosis

Mouse kidneys with a functional *Has1/Has3* double knockout (HAS1/3^-/-^) were investigated both histologically and through RT-qPCR for markers of fibrosis. *Has2* knockout mice demonstrate embryonic lethality and hence it was not possible to explore these. Naïve *Has1/3*^-/-^ mice had normal levels of serum creatinine and normal renal histology. However, they demonstrated lower levels of E-cadherin (Figure-8A&8B). They also demonstrated areas of interstitial α-SMA^+^ myofibroblast staining (Figure-8C), although the levels of α-SMA mRNA gene expression of the whole kidney did not show any statistical difference compared to wild-type (Figure-8D). Moreover, *Has1/3^-/-^* mice demonstrated small pockets of stromal HA staining in the renal cortex, which were absent in wild-type mice (Figure-8E) suggesting aberrant HAS2-driven HA synthesis in the renal cortex.

**Figure 8.**
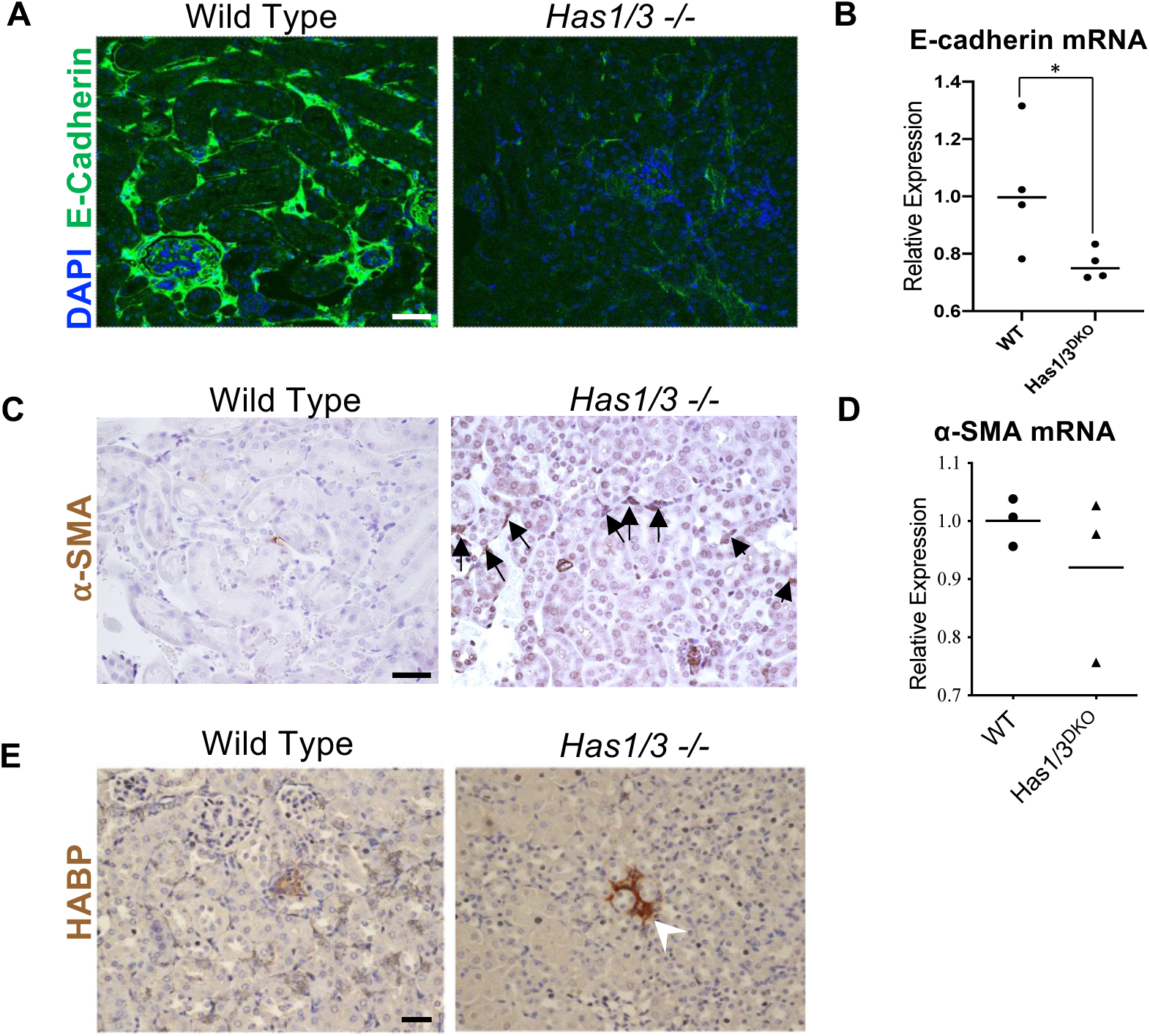
Mouse kidneys that only express the HAS2 isoenzyme demonstrate altered staining patterns of eCadherin, α-SMA and HA. Kidneys from 12-week-old C57BL6 Wild Type (n=3) and HAS1/3 double knockout (n=3) mice were harvested and shipped by Professor A. Plaas (Rush University, Chicago IL). Kidney specimens in RNAlater were homogenised and RNA was extracted from whole tissue for RT-qPCR analysis. Kidneys embedded in paraffin were sectioned, deparaffinised and histological sections prepared for immunofluorescence/immunohistology staining. Expression of (A, B) E-cadherin and (C, D) α-SMA were assessed by histology and QPCR. Expression of (E) HABP was assessed by immunohistology. Black arrows denote areas of increased α-SMA staining. White arrowhead denotes small pocket of interstitial HA staining. QPCR data are presented as dot plots representative of 3 separate experiments. \**P*<0.05; \*\**P*<0.01; \*\*\**P*<0.001. Scale bars 20 μm, Original Magnification x200.

### Alignment of HAS genomic DNA sequences reveals a conserved noncoding region (CNR) of upstream sequence shared by HAS1 and HAS2

Analysis of 3kb of sequence upstream of the transcription start sites of HAS1, HAS2 and HAS3 revealed a single CNR with a score >70% (supplementary data Figure-3A). The CNR was shared between HAS1 and HAS2 and spanned 165bp, with 72.1% sequence identity (supplementary data Figure-3B). No conserved sequence was identified as shared between HAS1 or HAS2 and HAS3.

To investigate the evolutionary history of the CNR we reconstructed branch lengths from a midpoint rooted tree using maximum likelihood as implemented by TreeFinder and 1,000 bootstrap replicates^39^. This reconstruction suggested that HAS2 and HAS3 are more closely related to each other than either is to HAS1, and that the CNR was lost in the HAS3 lineage (supplementary data Figure-3C).

## DISCUSSION

HA is a non-sulphated glycosaminoglycan, which is an important structural component of extracellular matrix. HA is ubiquitously expressed and interacts directly with cells regulating processes including cell-cell adhesion and crosstalk, migration, proliferation, and differentiation^40–44^. HA is therefore implicated in influencing numerous biological processes and has a vital role during mammalian organogenesis and development. Therefore, it is no surprise that dysregulated HA synthesis, turnover and binding interactions contributes to a multitude of disease states, including atherosclerosis, chronic inflammation, cancer and fibrosis^45–48^. In this study we demonstrate that during embryonic kidney development there is abundant HA in the stroma of the renal cortex. In the mature healthy kidney cortex, HA is absent but is rapidly re-expressed in the interstitial stroma following ischaemic injury. Furthermore, in mature kidneys protected from ischaemic damage, there is attenuated interstitial HA in the stroma. These collective findings suggest that renal HA may have dual and conflicting roles: stromal HA potentially plays an important role in nephrogenesis and in tubular regeneration following injury; however, persistence of stromal HA following injury promotes progressive fibrosis leading to nephron destruction rather than regeneration. Thus, understanding differences in HA generation in development *versus* injury may provide clues towards HA modification to promote regeneration rather than fibrosis.

HA is synthesized by the three HA Synthase iso-enzymes (HAS1, HAS2, HAS3). The principal cell-surface HA receptor is CD44, but HA can also bind to a multitude of HA binding proteins (termed Hyaladherins). The biological functions of HA are determined by its interactions with these hyaladherins, as well as its size and manner of synthesis and assembly within tissues. Thus, the biology of HA is complex, and HA can have diverse and sometimes opposing functions depending on these factors. Deletion of *Has2* in mouse embryos results in growth retardation evident by embryonic day-9 and the mice die at mid-gestation (before metanephric kidney development and ureteric bud outgrowth)^34^. By contrast, mice lacking one or both of the *Has1* and *Has3* genes are viable and exhibit no obvious phenotype^49, 50^. Our previous cell studies outlined the pathway via which HA regulates differentiation to an α-SMA^+^ myofibroblast phenotype, outlining the importance of a HAS2-generated cell-surface “HA coat” tethered to CD44 complexed with CD147 within the cell membrane^51^. This promoted MAPK/ERK signalling, which alongside SMAD signalling was critical for differentiation-to and maintenance-of the pro-fibrotic α-SMA^+^ myofibroblast phenotype^23, 27, 37, 52^. Our studies also showed that cytoplasmic internalisation and degradation of the HA coat can both prevent and reverse α-SMA^+^ differentiation, indicating that continued presence of a HA matrix can promote pro-fibrotic effects at a cellular level^25, 53^.

This study builds on our previous cell studies, confirming that increased HAS2 and interstitial HA in the renal cortex *in vivo* correlates with increased interstitial α-SMA^+^ myofibroblasts and promotes pro-fibrotic outcomes. We also confirm that fibroblasts with augmented HAS2 expression are pro-contractile and display enhanced expression of pro-fibrotic markers including EDA fibronectin and collagen type-I, whilst having attenuated expression of protective proteins such as FAP. Interestingly, genetically modified mice only capable of HAS2-driven HA generation appear to be “primed” towards developing renal injury with increased interstitial pockets of α-SMA and HA, even without renal injury. These findings are in keeping with studies looking at the fibrotic effects of HAS2 in models of skin and lung fibrosis^54–57^.

In contrast to HAS2, the literature on HAS1 and its role in HA generation, particularly in the kidneys is limited. HAS1 has received the least attention with its biological effects on cellular behaviour and association with disease states remaining largely unknown, and why vertebrates have three distinct HAS isoforms that are conserved through evolution remains an enigma. Early studies suggest that HAS1 has the lowest activity of all the HAS enzymes and its role in biology may be limited to providing compensatory HA synthesis for HAS2 and HAS3, which are depicted as the most active and biologically crucial HAS enzymes^59^. However, our report indicates that HAS1 has distinct biological functions from HAS2 in the kidney and in stromal cells. HAS1 is widely expressed and present during embryonic kidney development, particularly where HA is abundant in the stroma around developing tubules and areas of new capillary formation. In healthy mature kidneys, HAS1 has limited expression in the apical aspect of distal tubular epithelial cells, whilst in IRI-damaged kidneys, HAS1 expression is attenuated, disordered, and notably absent from areas of increased HA expression where HAS2 predominates. This indicates that HAS1 is not responsible for stromal HA synthesis during renal injury. However, in our protective renal model HAS1 is enhanced in the vasculature, in the distal tubular epithelial cells and notably in a population of interstitial cells that are distinct from α-SMA^+^ myofibroblasts. In cells studies, *HAS1* expression was markedly enhanced following stimulation with the renoprotective cytokine bone morphogenic protein-7 (BMP7). HAS1^+^ fibroblasts also had attenuated expression of pro-fibrotic markers (*ACTA2, EDA-FN, COL1A1*) and were less contractile, but demonstrated increased *FAP* expression and were pro-migratory. FAP is known to mediate collagen clearance in models of lung disease, leading to resolution of fibrous scarring^38^. Thus, the increased expression of HAS1 in IPC, the BMP7-mediated promotion of *HAS1* and the augmented FAP expression in HAS1^+^ fibroblasts (alongside attenuated expression of pro-fibrotic markers) collectively indicate a protective role for HAS1 in the kidney. Single-cell sequencing studies from lung tissues supports distinct roles for HAS1^+^ and HAS2^+^ stromal cells with a recent study indicating in lung fibrosis there is a stromal cell type marked by Has1 and this is separate from the α-SMA^+^ myofibroblast cluster that is associated with Has2^16^.

Fibroblasts can specialise in different tissues and have distinct phenotypes depending on their site of isolation. Advances in lineage tracing and single-cell transcriptional profiling techniques have also revealed impressive diversity amongst fibroblasts in a range of organ systems including the skin, lung, kidney and heart. In this report, not only do we demonstrate fibroblast subtypes as a useful tool to study stromal cell responses, but also demonstrate that modulation of HA, and in particular HAS isoenzymes, as an important regulator of fibroblast phenotypic heterogeneity; thereby highlighting HA as an important area of further study in the growing field of stromal cell heterogeneity.

## AUTHOR CONTRIBUTIONS

Irina Grigorieva, Adam Midgley and Aled Williams were involved in the study design, conducting experiments, data collection, and analysis or interpretation. Charlotte Brown and Usman Khalid performed the animal experimentation. Dominic Manetta-Jones undertook a proportion of the histological analysis. Gilda Chavez was involved as a blinded pathologist in analysis of the immunohistology of the mice and rat kidney sections. Anna Plaas and Deva Chan provided the kidney tissues from the Has1 knockout and Has1/3 double knockout mice as well as providing intellectual input into the HAS1 related data. Robert Steadman, Rafael Chavez, Timothy Bowen and Usman Khalid provided intellectual input. Soma Meran directed the study conception and design, provided the intellectual input, was involved in data analysis and prepared the manuscript drafts and revisions. All authors approved the final manuscript.

## ACKNOWLEDGEMENTS

The authors would like to thank Professor Ellen Pure and Dr Jamie Monslow from the University of Pennsylvania for the use of FAP primers for qPCR analysis.

## Supplementary Material to

**Supplemental Table 1:**
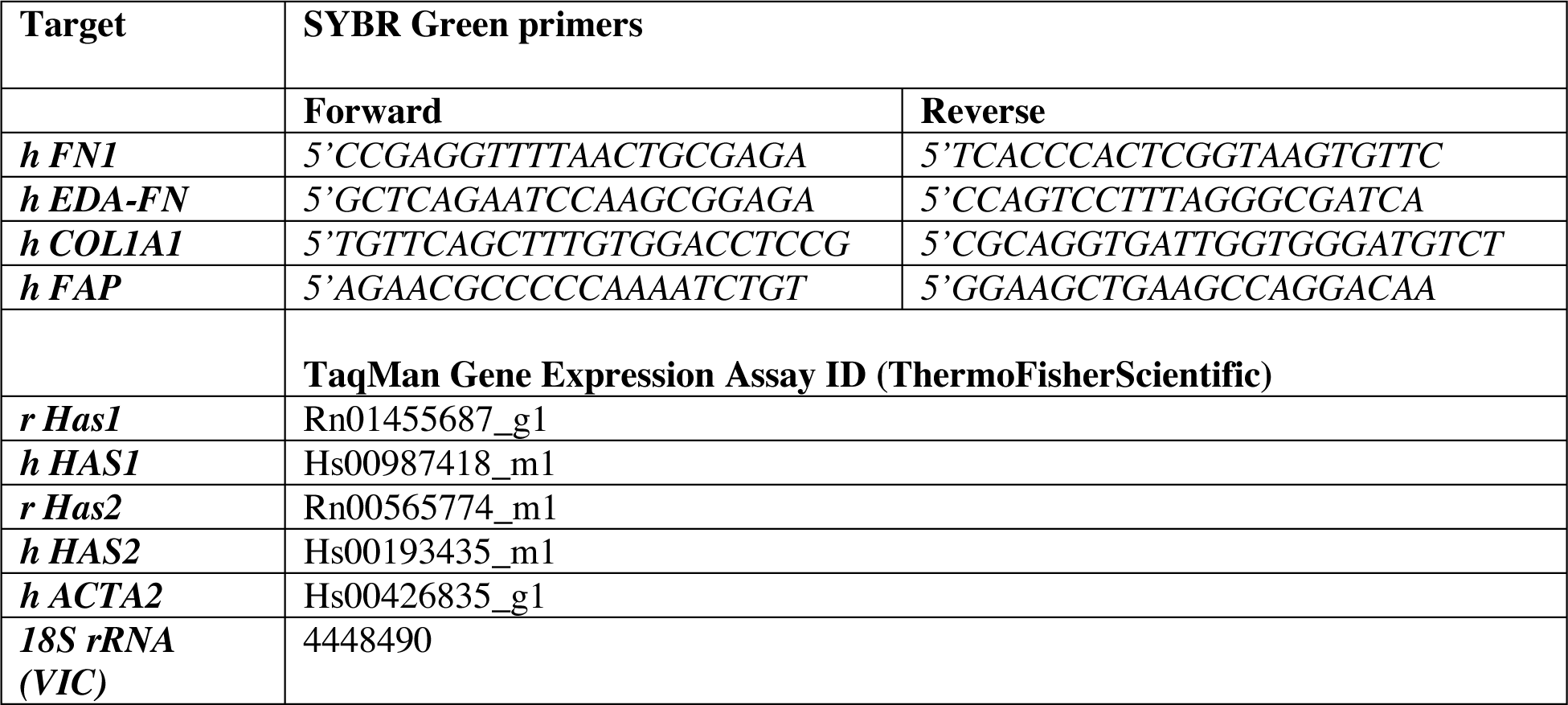
Primer Sequences for Real-Time PCR.

**Supplemental Table 2:** Primary antibodies.

| Antigen | Host | Source | Catalogue number | Dilution |
| --- | --- | --- | --- | --- |
| **α-smooth muscle actin (α-SMA)** | mouse | Thermo Fisher Scientific | MA5-11547 | 1:500 |
| **HAS1** | rabbit | Abcam | Ab 198846 | 1:200 |
| **HAS2** | mouse | SantaCruz | SC514737 | 1:200 |
| **E-cadherin** | rabbit | Sigma Prestige | HPA004812 | 1:500 |

**Supplemental Figure 1:**
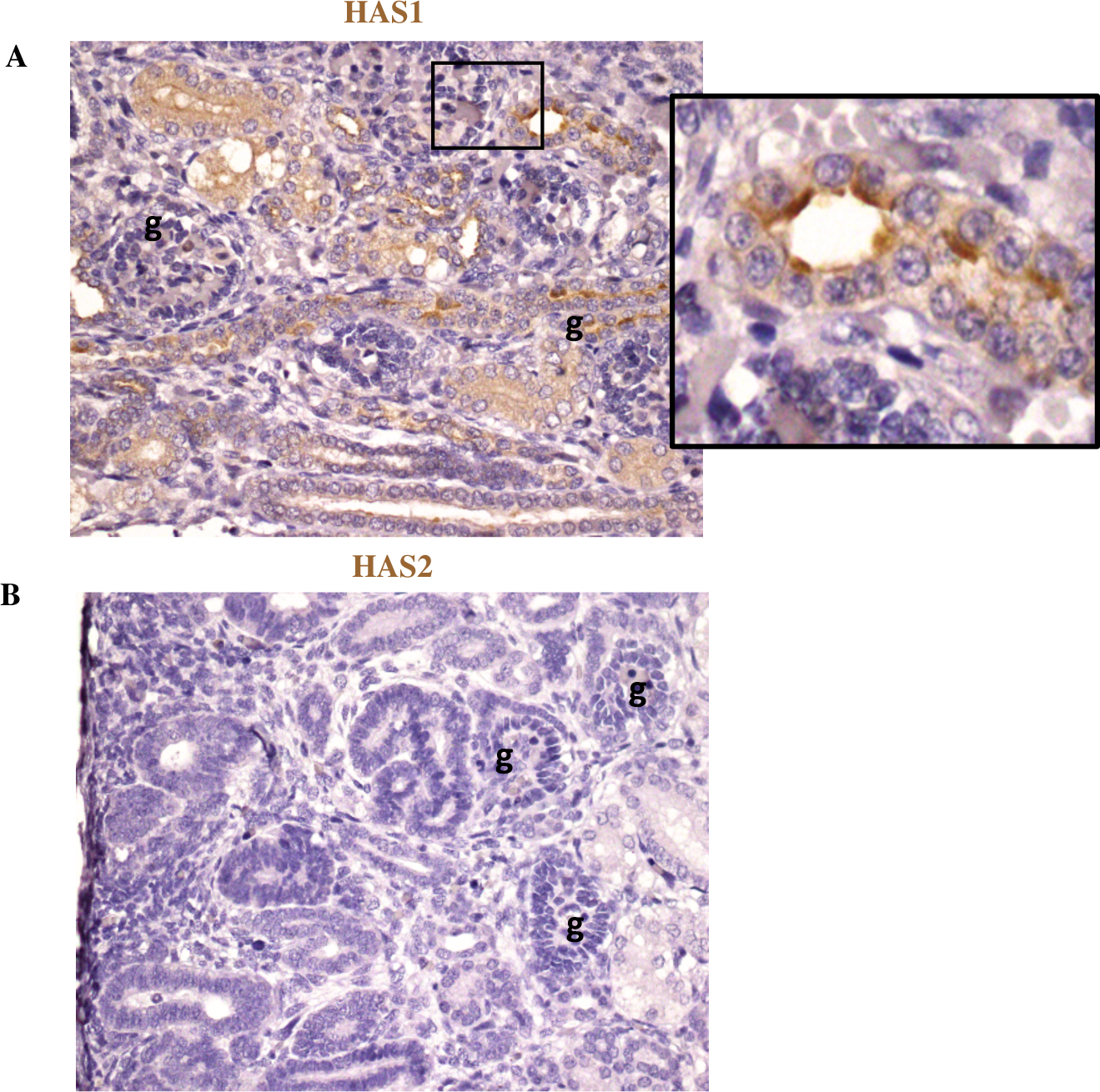
E15.5 mice kidneys stained for HAS1 or HAS2 using immunohistochemistry. Localisation of HAS1 and HAS2 iso-enzymes in developing mice kidneys. *Embryos* were collected with the day of the vaginal plug designated as embryonic day (E) 0.5 and staged by morphologic criteria, which included somite number and eye and limb morphology. Kidneys were dissected from E15.5 embryos. Immunohistochemical detection of (A) HAS1 and (B) HAS2 demonstrating presence of HAS1 and absence of HAS2 in developing mice kidneys at E15.5.

**Supplemental Figure 2:**
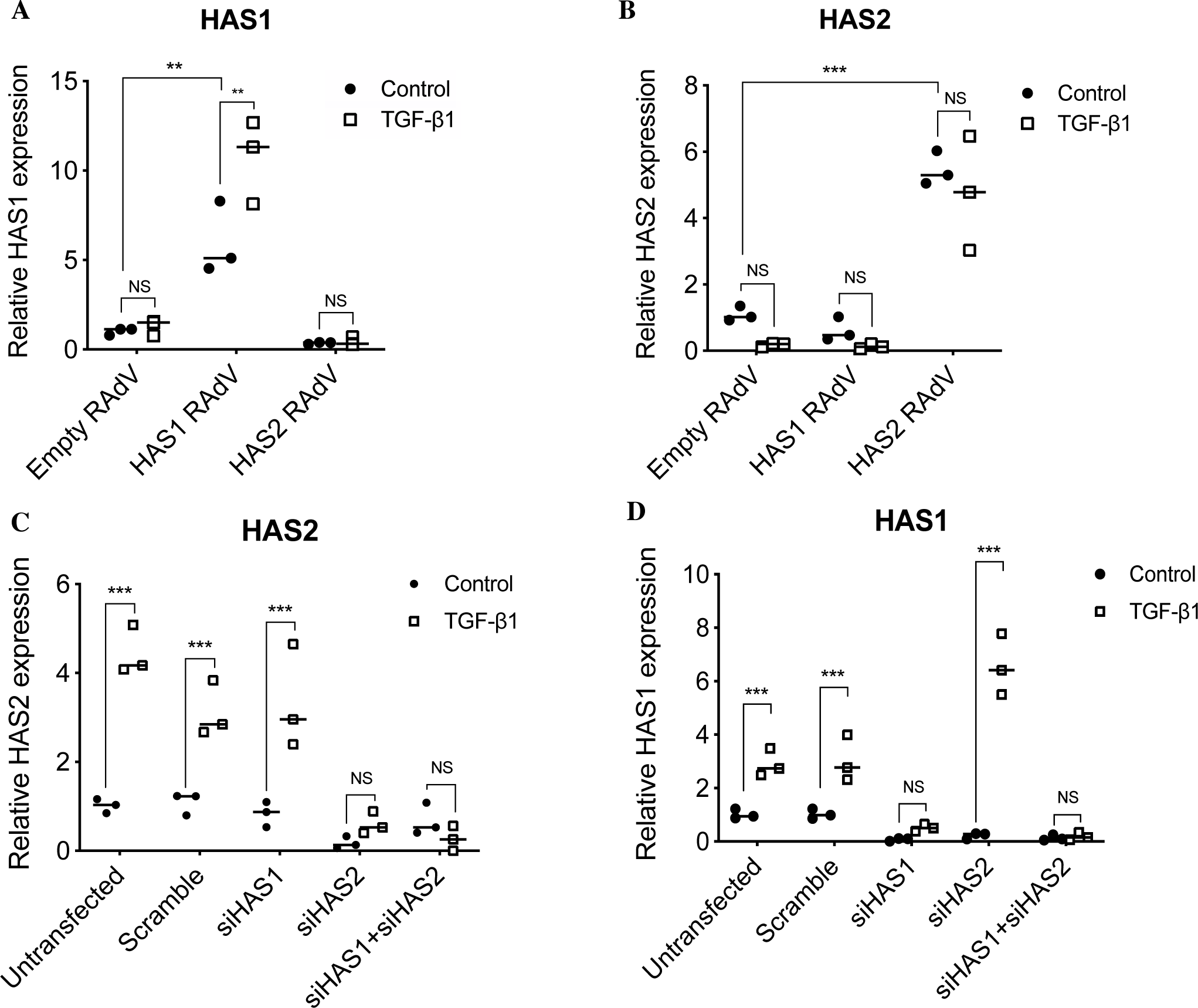
Confirmation of HAS isoenzyme over-expression and knockdown in fibroblasts. Confirmation of HAS isoenzyme over-expression and knockdown. (A, B) Empty recombinant adenoviral vector (RAdV) and HAS1 or HAS2 RAdVs were used to infect oral mucosal fibroblasts in the absence (●) or presence (□) of 10 ng/ml TGF-β1 for 72 hours. RT-qPCR was used to assess mRNA expression of HAS1 and HAS2 to confirm over-expression. (C, D) Dermal fibroblasts were transfected with scrambled siRNA or siRNA targeting HAS1, HAS2 or both and incubated in the absence (●) or presence (□) of 10ng/ml TGF-β1 for 72 hours. RT-qPCR was used to assess mRNA expression of Fibronectin (FN1), EDA-splice variant fibronectin (EDA-FN) and collagen-1 (COL1A1). Data are presented as dot plots representative of 3 separate experiments. \**P*<0.05; \*\**P*<0.01; \*\*\**P*<0.001.

**Supplementary Figure 3.**
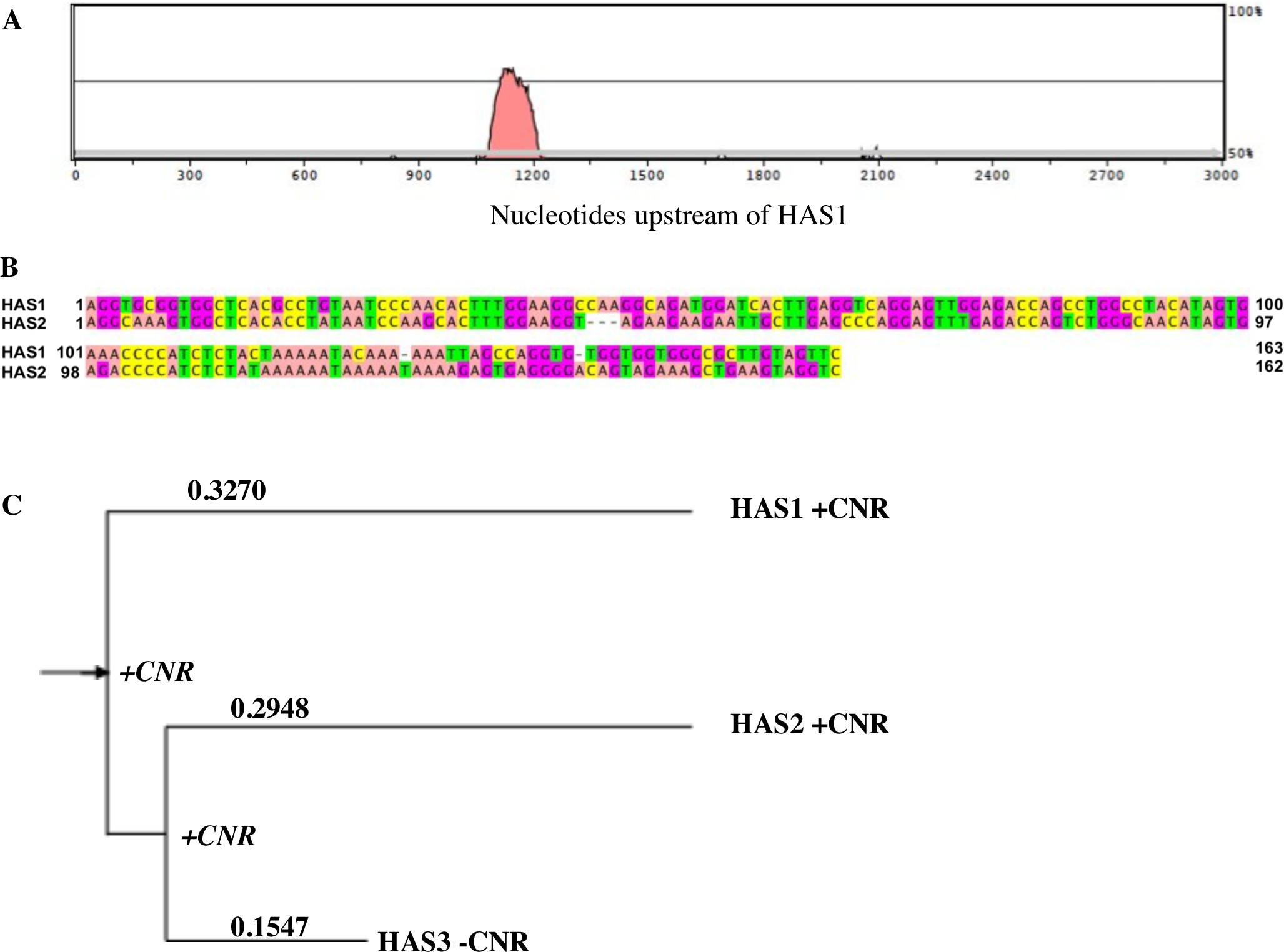
Human HAS gene promoter sequence alignment and phylogenetic HAS analysis. (A) VISTA output for a comparison of 3 kb upstream regions from HAS1 (transcript HAS1-001; Ensembl genomic coordinates Chromosome 19: 52,216,365-52,227,247 reverse strand) and HAS2 (transcript HAS2-001; Chromosome 8: 122,624,356-122,653,630 reverse strand). Sequences were extracted from Ensembl release 70, which uses the GRCh37.p10 assembly. The red-shaded region represents a sequence >70% conserved between HAS1 and HAS2 and is shown with nucleotide positions relative to the HAS1 transcription start site at 3001. (C) LAGAN alignment of the conserved region identified by VISTA, visualised using JALVIEW. (D) Phylogenetic analysis of human HAS gene cDNA sequences extracted from the Ensembl database. Sequences analysed were HAS1-001 (ENST00000540069.2; labelled here as HAS1), HAS2-001 (ENST00000303924.4; HAS2) and HAS3-001 (ENST00000306560.1; HAS3). Numbers on the branches are branch lengths (sequence differences) from the midpoint of the tree, which is marked with an arrow. The presence or absence of the CNR is indicated by “+CNR” for presence and “-CNR” for absence. Inferred presence of the CNR at branch points is indicated in italics.

